# Changes in genome organization of parasite-specific gene families during the *Plasmodium* transmission stages

**DOI:** 10.1101/242123

**Authors:** Evelien M. Bunnik, Kate B. Cook, Nelle Varoquaux, Gayani Batugedara, Jacques Prudhomme, Lirong Shi, Chiara Andolina, Leila S. Ross, Declan Brady, David A. Fidock, Francois Nosten, Rita Tewari, Photini Sinnis, Ferhat Ay, Jean-Philippe Vert, William Stafford Noble, Karine G. Le Roch

## Abstract

The development of malaria parasites throughout their various life cycle stages is controlled by coordinated changes in gene expression. We previously showed that the three-dimensional organization of the *P. falciparum* genome is strongly associated with gene expression during its replication cycle inside red blood cells. Here, we analyzed genome organization in the *P. falciparum* and *P. vivax* transmission stages. Major changes occurred in the localization and interactions of genes involved in pathogenesis and immune evasion, erythrocyte and liver cell invasion, sexual differentiation and master regulation of gene expression. In addition, we observed reorganization of subtelomeric heterochromatin around genes involved in host cell remodeling. Depletion of heterochromatin protein 1 (PfHP1) resulted in loss of interactions between virulence genes, confirming that PfHP1 is essential for maintenance of the repressive center. Overall, our results suggest that the three-dimensional genome structure is strongly connected with transcriptional activity of specific gene families throughout the life cycle of human malaria parasites.

## INTRODUCTION

With an estimated 445,000 deaths per year, malaria is still one of the most deadly infectious diseases, mostly targeting young children in sub-Saharan Africa^1^. The disease is caused by one of five parasites of the *Plasmodium* species, of which *P. falciparum* is the most common and deadly variant. *P. vivax* is also responsible for significant disease, mostly in Southeast Asia^1^.

*Plasmodium* parasites have complex life cycles that involve a human host and a mosquito vector. Infection in humans starts when an infected female mosquito takes a blood meal and transmits parasites that are present in the form of sporozoites in her salivary glands. These sporozoites are inoculated into the skin, travel to the liver and establish an infection in hepatocytes. Over a period of several days, the parasite replicates and eventually releases thousands of merozoites into the bloodstream. In addition, *P. vivax* can survive in the liver for weeks or years in dormant forms called hypnozoites, which can be reactivated and cause malaria relapses. Merozoites that emerge from the liver start a 48-hour replication cycle in red blood cells. During this asexual intraerythrocytic development cycle (IDC), the parasite progresses through three main developmental stages: ring, trophozoite, and schizont, to produce approximately 16 daughter parasites, which burst from the cell and invade new erythrocytes. During the IDC, the parasite can commit to differentiation into male and female gametocytes, which can be taken up by another mosquito. Inside the mosquito, the parasite undergoes sexual reproduction and further develops through several stages into the salivary gland sporozoites that can be transmitted to a new human host.

Understanding how the transitions between the various life cycle stages of the *Plasmodium* parasite are regulated remains an important goal in malaria research. Stage transitions are regulated by coordinated changes in gene expression, but it is still largely unknown how these changes in transcriptional profiles are controlled at the transcriptional level. Only a single family of ApiAP2 transcription factors (TFs) with 27 members has been identified, while approximately two-thirds of the TFs expected based on the size of the *Plasmodium* genome seem to be missing^2–4^. Several of these ApiAP2 TFs are involved in stage transitions, such as PfAP2-G, which is thought to be the main driver for gametocyte differentiation^5–7^. Our understanding of how these TFs are controlled and how various TFs may act together to form transcriptional networks is still very limited.

Over the past years, we have gained much insight into the role of epigenetics, chromatin structure and genome organization in gene regulation, mostly during the IDC of *P. falciparum*. It has become clear that the parasite genome is largely in an active, euchromatic state^8–11^, while members of several parasite-specific gene families involved in virulence *(var, rifin, stevor*, and *pfmc-2tm)*, erythrocyte remodeling *(phist, hyp, fikk*, and others) and solute transport *(clag3)* are organized into heterochromatin^9,12–15^. In particular, the family of *var* genes has received much attention, since these genes are extremely important for pathogenesis and immune escape. Out of a total of 60 *var* genes, only a single variant is expressed within an individual parasite, while the other 59 genes are tightly repressed by a combination of isolation into a perinuclear compartment, repressive histone marks, and repressive long non-coding RNAs^9,16–19^.

Previously, we assessed genome organization at the ring, trophozoite, and schizont stages of the IDC in *P. falciparum* using Hi-C experiments (chromosome conformation capture coupled with next-generation sequencing)^20^ and compared our findings to an earlier Hi-C study in ring-stage *P. falciparum*^21^. We observed a strong association between genome architecture and gene expression, suggesting that the three-dimensional organization of the genome is very important for gene regulation. Here, we analyzed the genome organization in the transmission stages of the *P. falciparum* life cycle (gametocytes and sporozoites), as well as for *P. vivax* sporozoites, and present a comparative analysis of genome organization throughout the different life cycle stages. Finally, to meaningfully compare changes in genome organization throughout the *Plasmodium* life cycle, we developed a novel statistical test to detect loci that differ significantly in their intrachromosomal contact count numbers between pairs of stages. This methodology can be directly applied to pairs of Hi-C data sets from any organism to assess chromosome dynamics.

## RESULTS

### Capturing the genome conformation of *Plasmodium* transmission stages

To complement our previous study describing the genome architecture of *P. falciparum* during the intraerythrocytic developmental cycle (IDC)^20^, we performed Hi-C experiments on three additional stages of the *P. falciparum* life cycle: early gametocytes (stage II/III), late gametocytes (stage IV/V), and salivary gland sporozoites (**Figure 1A**, **Supplemental Table S1**, and **Supplemental Fig. S1**) using the tethered conformation capture methodology. In addition, to evaluate similarities and differences in genome organization between the highly pathogenic *P. falciparum* and the less virulent *P. vivax*, we generated Hi-C data for *P. vivax* salivary gland sporozoites. For each stage, we obtained high-quality data, evidenced by a log-linear relationship between contact probability and genomic distance (**Supplemental Fig. S2A**), as well as interchromosomal contact probability (ICP) and percentage of long-range contacts (PLRC) values in agreement with previous studies (**Supplemental Table S2**). Biological replicates of both *P. falciparum* and *P. vivax* sporozoite stage parasites showed a high degree of similarity (**Supplemental Fig. S2B-C**), demonstrating the robustness of our methodology. The data from these replicates were combined to obtain higher resolution for subsequent analyses.

**Figure 1:**
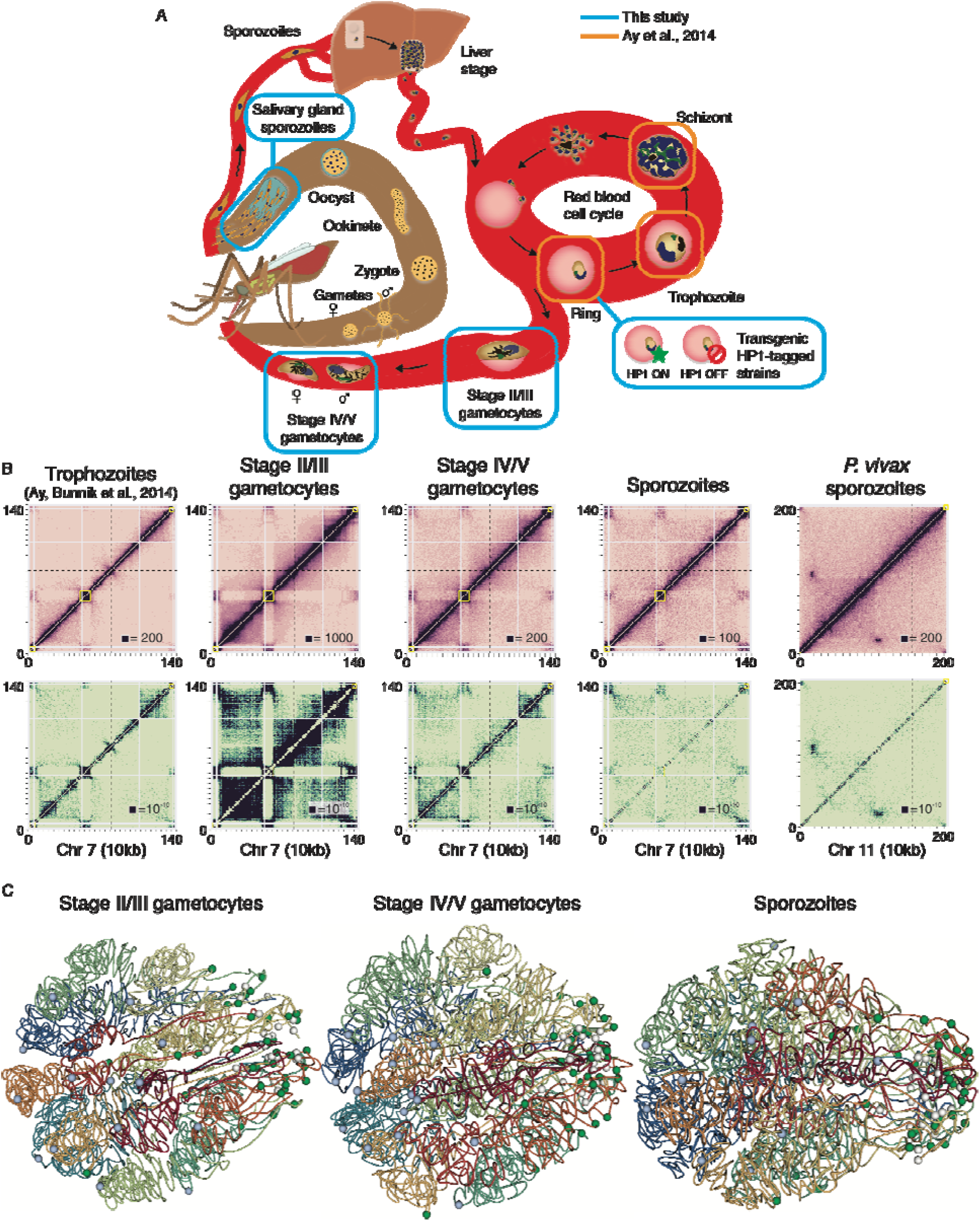
Genome organization in *Plasmodium* parasites. **(A)** Schematic overview of the parasite life cycle, with the samples generated in this study highlighted in blue and stages available from a previous study^20^ shown in orange. **(B)** ICE-normalized contact count matrices (top row) and fit-hi-c p-value matrices (bottom row) at 10 kb resolution of chromosome 7 for *P. falciparum* stages and chromosome 11 for *P. vivax* sporozoites. The boxed value indicates the maximum contact count (top row) or minimum P-value (bottom row). In all other figures comparing different stages, the contact counts were subsampled to the same total. Virulence clusters are indicated by yellow boxes, the centromere location by a dashed black line, and unmappable regions by grey in these and all other heatmaps. **(C)** Models of the consensus three-dimensional organization of the *P. falciparum* genome in stage II/III gametocytes, stage IV/V gametocytes and salivary gland sporozoites, with light blue spheres indicating centromeres, white spheres indicating telomeres and green spheres indicating the location of virulence gene clusters.

Next, the observed intrachromosomal and interchromosomal contacts were aggregated into contact count matrices at 10-kb resolution and were normalized using the ICE method to correct for experimental and technical biases^22^ (**Figure 1B**, **Supplemental File S1**, and **Supplemental File S2**). In addition, we identified significant contacts using fit-hi-c, which controls for the propensity of adjacent loci to have more contacts and calculates a p-value reflecting the probability that the number of contacts in that bin is larger than expected by chance^23^. We inferred a consensus 3D genome structure for each of the transmission stages and the three IDC stages using Pastis^24^ (**Figure 1C**). Pastis uses a maximum likelihood approach based on modeling contacts using a negative binomial model to account for the observed overdispersion. This method generates more accurate and robust structures when compared to the optimization-based approach we used previously^20^. The stability of these consensus structures was assessed by generating 5,000 possible structures from varying initial starting points. A principal component analysis showed strong clustering of structures from the same stage, and clear separation between structures of different stages, except for early and late gametocytes (**Supplemental Fig. S2D**). These results indicate that the genome organization of early and late gametocytes is similar, while there are distinct differences between all other stages that are captured using a single representative structure for each stage.

### Universal and stage-specific features of genome organization

From the contact count matrices and the consensus 3D structures, it became apparent that large-scale features of genome organization at the gametocyte stage were comparable to those of the IDC stages, including colocalization of centromeres and clustering of telomeres and virulence genes (all p-values = 0.0, Witten-Noble colocalization test^25^). However, we observed significant intrachromosomal rearrangements, including increased interactions among virulence genes and exported proteins, repression of invasion genes, change in organization of ribosomal DNA genes, as well as the formation of large domains on chromosome 14 in close proximity to a female gametocyte-specific *pfap2* transcription factor locus. These changes will be addressed in more detail in the next sections. At the sporozoite stage of *P. falciparum*, the clustering of telomeres was conserved, but the colocalization of the centromeres was completely lost (p-value = 0.49, **Figure 1C**, rightmost panel, and **Supplemental Fig. S3A**). In contrast, in *P. vivax* sporozoites, the centromeres colocalized significantly (p-value = 0.0), but these interactions were only observed at the centromere itself and did not involve any of the surrounding regions (**Supplemental Fig. S3B**). To the best of our knowledge, IFA on nuclear proteins and DNA-FISH have never been successfully performed in the sporozoite stage. While our result will need to be validated by an independent approach, our Hi-C experiment is so far the only successful technique that has been able to monitor a reduction in centromere clustering in the sporozoite stage. To better visualize the large-scale differences between the various stages of the *P. falciparum* life cycle, we generated an animation of the changes in genome organization during the stage transitions, which highlights that chromosomes undergo dramatic rearrangement in sporozoites as compared to the blood stages (Supplemental Movie S1).

To systematically identify changes in genome conformation between the various stages, we designed a statistical test that analyzes differences in the number of intrachromosomal contacts for each 10-kb bin in the normalized contact count matrices between pairs of stages. Briefly, we modeled each bin as a negative binomial random variable, and we estimated the relationship between the mean and variance by grouping pairs of loci that are separated by the same linear genomic distance. We then used an exact negative binomial test, similar in form to a Fisher’s exact test and previously applied to differential expression analysis^26,27^ to calculate a p-value (see **Computational Methods** in **Supplemental Information**). Contact count matrices were subsampled to account for differences in interaction counts measured at the various stages and only intrachromosomal contacts for which the sum of the contacts was above the 80% percentile were tested. The resulting fold-change values were filtered after FDR estimation to only include loci that show a 2-fold or larger difference in contacts, after normalizing for the effect of genomic distance, with a false discovery rate of less than 1% (see **Computational Methods** in **Supplemental Information**; **Supplemental Table S3**). As expected, interactions at the sporozoite stages were most different from those in other stages of the life cycle. Several chromosomes showed large rearrangements towards the chromosome ends, for example the right arm of chromosome 4 (**Supplemental Fig. S4**), involving gene families coding for exported proteins involved in virulence and erythrocyte remodeling. The fold-change heatmaps (accessible at http://noble.gs.washington.edu/proj/plasmo3dsexualstages/), the contact count heatmaps, and the confidence score heatmaps showed many additional differences in chromosome conformation during stage transitions, as described in more detail below. The changes in intrachromosomal and interchromosomal interactions in sporozoite and gametocyte stages were observed in two distinct field isolates or laboratory strains respectively, suggesting that genetic variations did not influence our results. Furthermore, the Hi-C methodology has recently been used to detect translocations in genomes and to correct genome assemblies based on Illumina and/or PacBio sequencing in many organisms, including *Plasmodium knowlesi, Arabidopsis thaliana* and *Aedes aegyptF*^8^'^34^. It is therefore unlikely that the changes that we observed between life cycle stages are artifacts caused by genomic recombination during *in vitro* parasite culture, since none of the interchromosomal heatmaps (**Supplemental Fig. S3A**) showed any evidence of such recombination events. To further validate our results, we have introduced interchromosomal and intrachromosomal translocations in the *P. vivax* genome to visualize the aberrant patterns that such recombination events would produce (**Supplemental Fig. S5**). In addition, we scanned our samples using a recently published metric developed to detect genome assembly errors in Hi-C data^33^ and did not detect any signs of misassembly or translocations (**Supplemental Fig. S6**).

### *pfap2-g* dissociates from the repressive center during gametocytogenesis

Previous work has shown that knockdown of heterochromatin protein 1 (PfHP1) during the IDC results in activation of the gametocyte-specific transcription factor locus *pfap2-g* and an increased formation of gametocytes^35^. PfHP1 interacts with the repressive histone mark H3K9me3 on silenced *var* genes that are colocalized in a perinuclear heterochromatic compartment. Using DNA-FISH, we observed that the *pfap2-g* locus was located in close proximity to a subtelomeric *var* gene on chr8 in >90% of the cells observed (**Figure 2A** and **Supplemental Fig. S7A**), suggesting that *pfap2-g* is associated with the repressive cluster during the IDC. In agreement with this observation, the trophozoite and schizont stage fit-hi-c p-value heatmaps showed a significant interaction between *pfap2-g* and the nearest internal virulence gene cluster (**Figure 2B** and **Supplemental Fig. S8A**). The virulence cluster and *pfap2-g* locus each straddle two ten kilobase bins, and in trophozoites, each of the four possible pairwise interactions were significant (fit-hi-c q-value < 0.05). We performed virtual 4C at MboI restriction site resolution to demonstrate that this interaction was specific for *pfap2-g* and did not involve the nearby *pfap2* PF3D7_1222400 (**Supplemental Fig. S9**). In addition, *pfap2-g* significantly interacted with virulence clusters on chromosomes 6 and 8 in trophozoites (fit-hi-c q-value < 0.05; **Supplemental Table S4**). In stage II/III gametocytes, no significant interactions between *pfap2-g* and virulence clusters were observed (q-values = 1.0; **Figure 2B**; **Supplemental Table S4**; **Supplemental Table S5**), indicating that *pfap2-g* dissociates from the repressive cluster in the transition from the IDC to early gametocytes. Interactions between *pfap2-g* and virulence clusters were partially regained in the late gametocyte stage (**Supplemental Table S4**, **Supplemental Table S5**). Unfortunately, DNA-FISH experiments were unsuccessful for the gametocyte stage. An improved DNA-FISH methodology for the gametocyte stages will need to be developed to further confirm these results by an independent approach.

**Figure 2:**
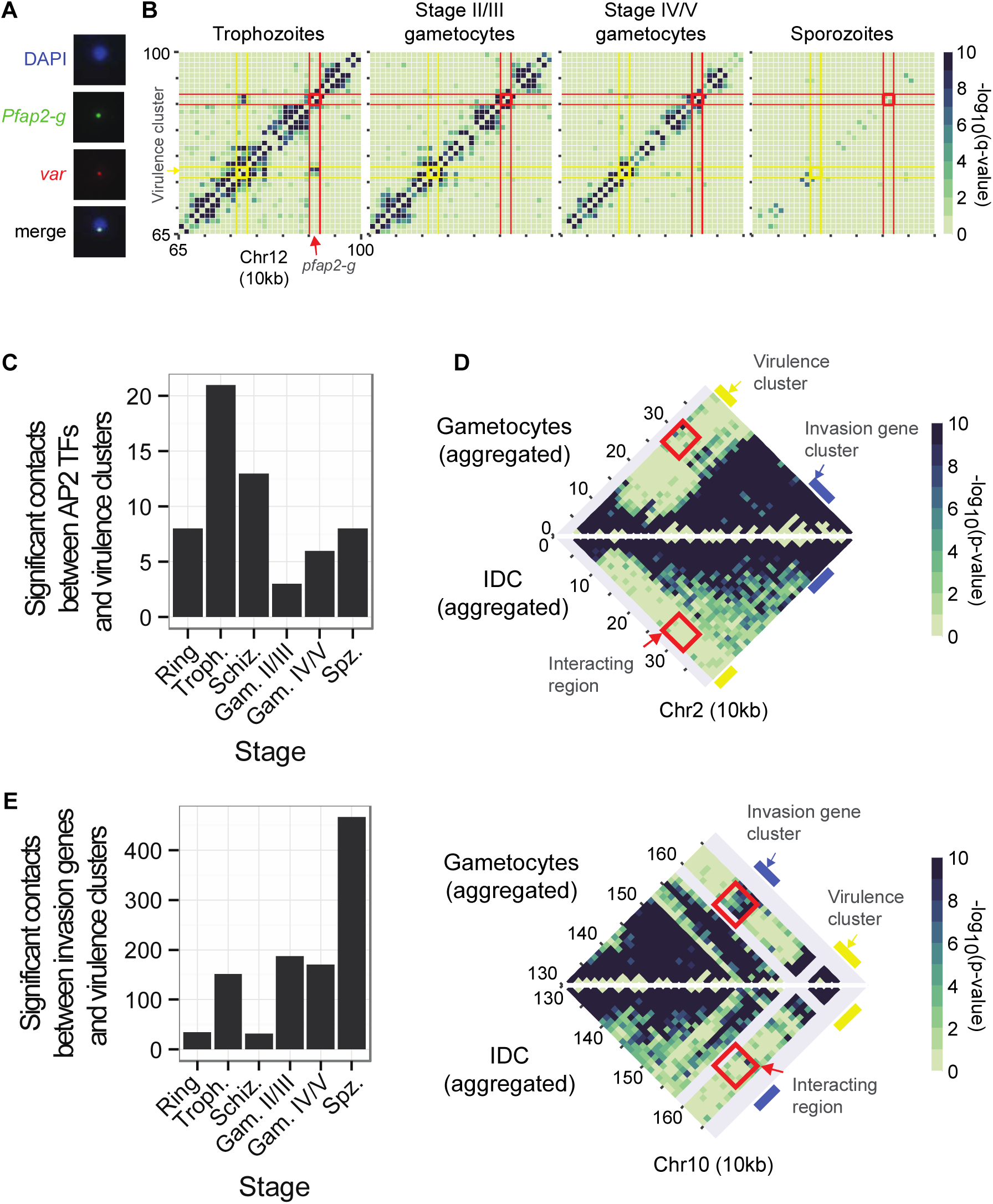
Changes in interaction of *pfap2* genes and invasion genes with the repressive center. **(A)** Colocalization of *pfap2-g* and *var* gene PF3D7_0800300 by DNA-FISH. Additional images are presented in Supplemental Fig. S6. **(B)** Dissociation of the gametocyte-specific transcription factor locus *pfap2-g* (red) from the nearby internal virulence gene cluster (yellow) in stage II/III gametocytes. **(C)** Overall reduced number of intrachromosomal and interchromosomal interactions between *pfap2* TF genes and virulence genes in gametocytes and sporozoites as compared to the IDC stages. **(D)** Invasion gene clusters (blue bar) on chromosomes 2 (top) and 10 (bottom) interact with subtelomeric virulence genes (yellow bar) in gametocytes, but not during the IDC. In each plot, the top triangle shows the aggregated data of both gametocyte stages, while the bottom triangle shows the aggregated data from the three IDC stages. Bins that depict interactions between virulence genes and invasion genes are highlighted by a red box. E) Increased number of intrachromosomal and interchromosomal interactions between invasion genes and virulence genes in gametocytes and sporozoites as compared to the IDC stages.

Significant contacts with virulence gene clusters at any parasite stage were observed for a total of 9 *pfap2* loci (**Supplemental Table S5**). The dissociation of *pfap2* genes from the repressive center in other stages than the IDC seemed to be a general trend: fewer significant contacts (q-value < 0.05) between *pfap2* genes and virulence gene clusters were observed in gametocytes and sporozoites as compared to the IDC stages (p-value = 0.0027, one sided sign test; **Figure 2C** and **Supplemental Fig. S8B**). We used a resampling procedure (see **Computational Methods** in **Supplemental Information**) to compare this result to randomly-selected bins; the p-value was 0.021. Note that *pfap2* PF3D7_0420300 is located in close genomic proximity to a virulence cluster in chromosome 4. To ensure that the significant decrease in *pfap2* gene loci interacting with virulence clusters is not due to this gene, we repeated the p-value calculation without PF3D7_0420300 and confirmed that it remained significant (p-value = 1.192e-07). Such changes in interactions with virulence gene clusters between stages were not observed for an unrelated gene family (i.e. histone genes, data not shown). These results are indicative of an important connection between genome organization and the activity of *pfap2* genes that drive parasite life cycle progression.

### Invasion genes relocate to the repressive center during gametocytogenesis

Distinct changes were observed in chromosomes 2 and 10 at loci that harbor invasion genes (**Figure 2D**). These genes encode proteins that are expressed in merozoites and mediate attachment to and entry of red blood cells and include several merozoite surface proteins, S-antigen, and glutamate-rich protein. In contrast to several other invasion genes *(pfrh4*^36^*, clag*, and eba^37^, these genes are not known to undergo clonally variant expression, and we therefore consider their association with heterochromatin in gametocytes to be a different regulatory mechanism than the epigenetic mechanisms that control expression of *pfrh4, clag*, and *eba* during the IDC. In gametocytes, these loci showed strong interaction with the subtelomeric regions, while these contacts were not observed in the IDC. To quantify this observation, we assessed the number of significant contacts between invasion gene loci and virulence clusters in each life cycle stage. A larger number of significant contacts were observed between invasion genes and virulence clusters in the transmission stages than in IDC stages (p-value < 2.2e-16, sign test; **Figure 2E** and **Supplemental Fig. S8C**). The lack of interactions between invasion gene GLURP on chromosome 10 (PF3D7_1035300) and *var* gene PF3D7_0800300 during the IDC was confirmed by DNA-FISH (**Supplemental Fig. S7B**). As mentioned earlier, DNA-FISH experiments were unfortunately not successful for the gametocyte stage. Collectively, our data indicate that, similar to the *pfap2* gene family, the expression of invasion genes during the life cycle is correlated with association with or dissociation from repressive heterochromatin.

### Expansion of subtelomeric heterochromatin in gametocytes

To further study changes in chromatin organization during gametocytogenesis, we determined the distribution of repressive histone mark H3K9me3 in late ring/early trophozoites and stage IV/V gametocytes by performing ChIP-seq on two biological replicates for each stage (**Supplemental Fig. S10A** and **Supplemental Table S6**) that were combined for downstream analyses. We used two different commercially available anti-H3K9me3 antibodies to rule out that our ChIP-seq results were influenced by the antibody used. In trophozoites, H3K9me3 marking was restricted to subtelomeric regions, internal virulence gene clusters and a few additional loci (incuding *pfap2-g* and *dblmsp2)*, as previously described for both H3K9me3^9,11^ and PfHP1^38^ (**Figure 3A** and **Supplemental Fig. S10B-C**). These same regions were occupied by H3K9me3 in gametocytes. In addition, in several chromosomes, the subtelomeric heterochromatin marking expanded to more internally located genes in gametocytes (**Figure 3A-C** and **Supplemental Fig. S10B-C**). The expansion of heterochromatin at the chromosome ends was also visible in the contact count heatmaps as larger subtelomeric domains that showed strong intra-domain interactions and were depleted of interactions with the internal region of the chromosome (**Supplemental File S1**). While not all genes in these regions were marked by H3K9me3, a total of 79 genes showed increased H3K9me3 levels in gametocytes as compared to trophozoites, 61 of which were exported proteins (**Supplemental Table S6**).

**Figure 3:**
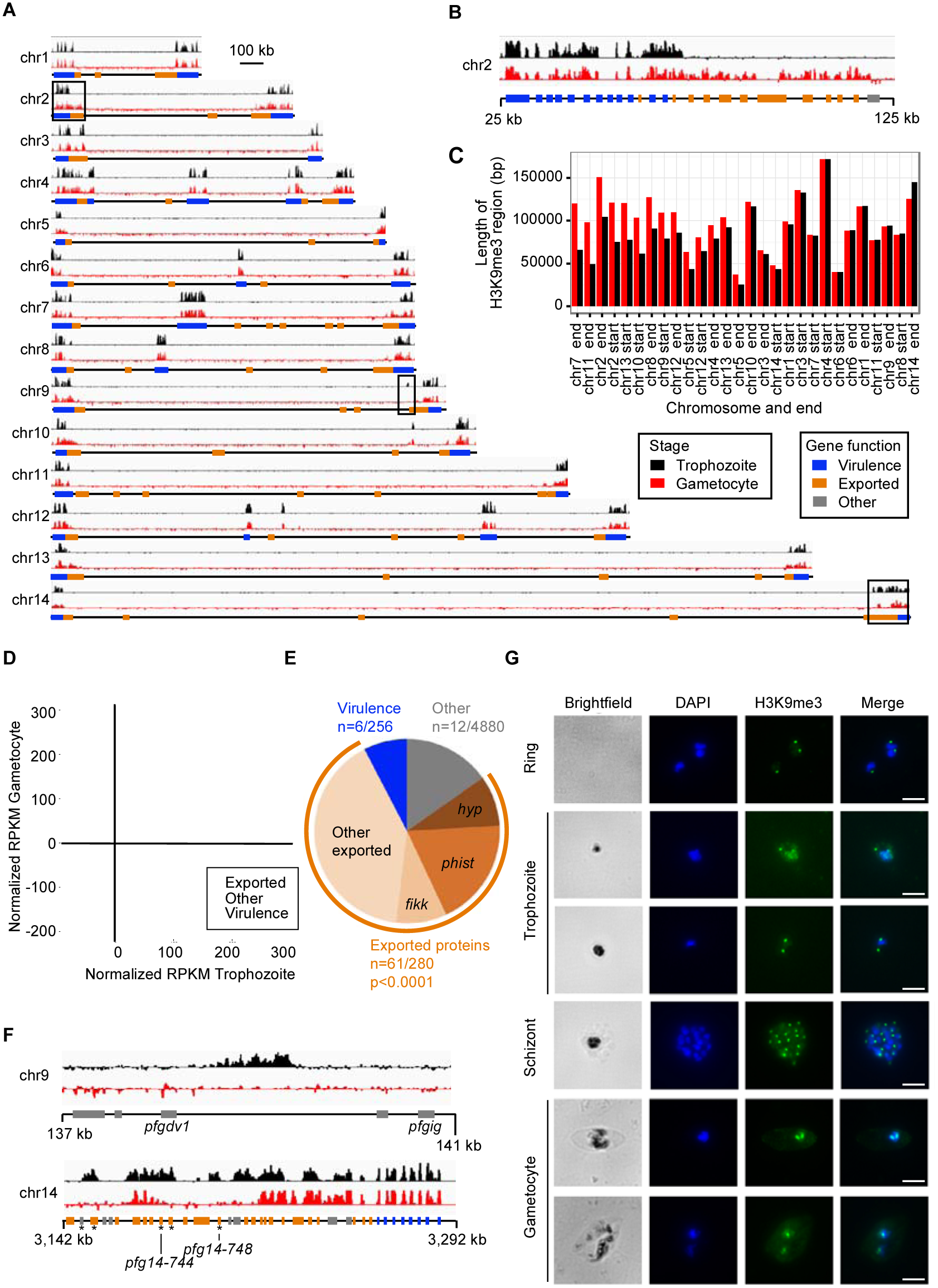
Silencing of genes encoding exported proteins in gametocytes through expansion of heterochromatin. **(A)** ChIP-seq analysis of genome-wide H3K9me3 localization in trophozoites (top tracks in black) and stage IV/V gametocytes (bottom tracks in red). Results of one representative biological replicate are shown for each stage. The regions depicted in panels B and F are indicated with black boxes. **(B)** Expansion of H3K9me3 heterochromatin in gametocytes as compared to trophozoites, predominantly to genes encoding exported proteins. **(C)** Length of each subtelomeric region in which the majority of genes is marked by H3K9me3, sorted by the difference in length between these regions in trophozoites and gametocytes. **(D)** H3K9me3 levels per gene at the trophozoite and gametocyte stages. **(E)** Enrichment of genes encoding for exported proteins among genes with increased levels of H3K9me3 in gametocytes (P-value from a two-tailed Fisher’s exact test). **(F)** Loss of H3K9me3 mark in gametocytes on chromosome 9 between gametocyte development genes *pfgdv1* and *pfgig*, as well as at gametocyte-specific genes encoding exported proteins on chromosome 14 (indicated with an asterisk). **(G)** Immunofluorescence analysis showing a single H3K9me3 focus in ring and schizont stages, and either one or two foci in gametocytes. Scale bar denotes 1 μm.

These genes included members of the *phist* (n=15 out of a total of 68), *hyp* (n=7 out of 34) and *fikk* (n=7 out of 19) families, as well as 32 other genes annotated as exported proteins or containing a PEXEL export motif (**Figure 3D-E**). Other members of gene families encoding exported proteins were marked with H3K9me3 in the IDC. However, 47 of these 61 genes have never been detected in a heterochromatic state in the IDC^9,11,38^. At two loci on chr9 and chr14, respectively, H3K9me3 was lost in gametocytes (**Figure 3F**). The locus on chr9 is between two genes known to be involved in gametocyte differentiation *(pfgdv1* and *pfgig)* and is deleted in various gametocyte-defective *P. falciparum* strains^39^. On chromosome 14, the genes that were not marked by H3K9me3 in gametocytes encode exported proteins that have been implicated in gametocytogenesis: PF3D7_1476600, PF3D7_1477300 *(Pfg14–744)*, PF3D7_1477400, and PF3D7_1477700 *(Pfg14–748)*^39^. This is the first data set that compares H3K9me3 marking between the IDC and gametocyte stages and implies that H3K9me3 and chromatin structure may play important roles in gene activation and silencing during gametocyte formation. To validate the 3D modeling and heterochromatin clustering, we performed immunofluorescence imaging against repressive histone mark H3K9me3. We identified a single nuclear H3K9me3 focus per nuclei in rings and schizonts (**Figure 3G** and **Supplemental Fig. S11**), corresponding to the single repressive center harboring all virulence genes as predicted by our 3D models. In trophozoites, the number of foci varied, in line with nuclear expansion^20,40^, and the progression of DNA replication that takes place at the end of this stage (**Figure 3G**). Gametocytes showed either one or two H3K9me3 foci that did not seem to be associated with a male or female phenotype. Further experiments will need to be performed to understand the biological relevance of the presence of this second focus in gametocytes.

### Formation of superdomains on a *P. falciparum* gametocyte chromosome

Chromosome 14 showed the formation of a strong domain boundary in both early and late gametocytes, which was not observed in any of the other life cycle stages (**Figure 4A** and **Supplemental File S1**). This separation of the chromosome into two superdomains is reminiscent of the bipartite structure of the inactivated X chromosome (Xi) in human, rhesus macaque, and mouse^41–44^. Zooming in on the boundary region of *P. falciparum* chromosome 14 showed a sharp transition at the MboI restriction site at nucleotide position 1,187,169 (**Figure 4B** and **Supplemental Fig. S12**). To demonstrate that this sharp boundary is not the result of a chromosomal translocation in the NF54 strain used for gametocyte isolations, we confirmed that the genomic region that spans the boundary region can be amplified by PCR and can be detected by Southern blot in both 3D7 and NF54 strains (**Supplemental Fig. S13**). In eukaryotic genomes, genes close to the domain boundary are often associated with higher levels of transcription^45,46^. The domain boundary is located inside or near PF3D7_1430100, which encodes serine/threonine protein phosphatase 2A activator (PTPA; **Supplemental Fig. S14A**).

**Figure 4:**
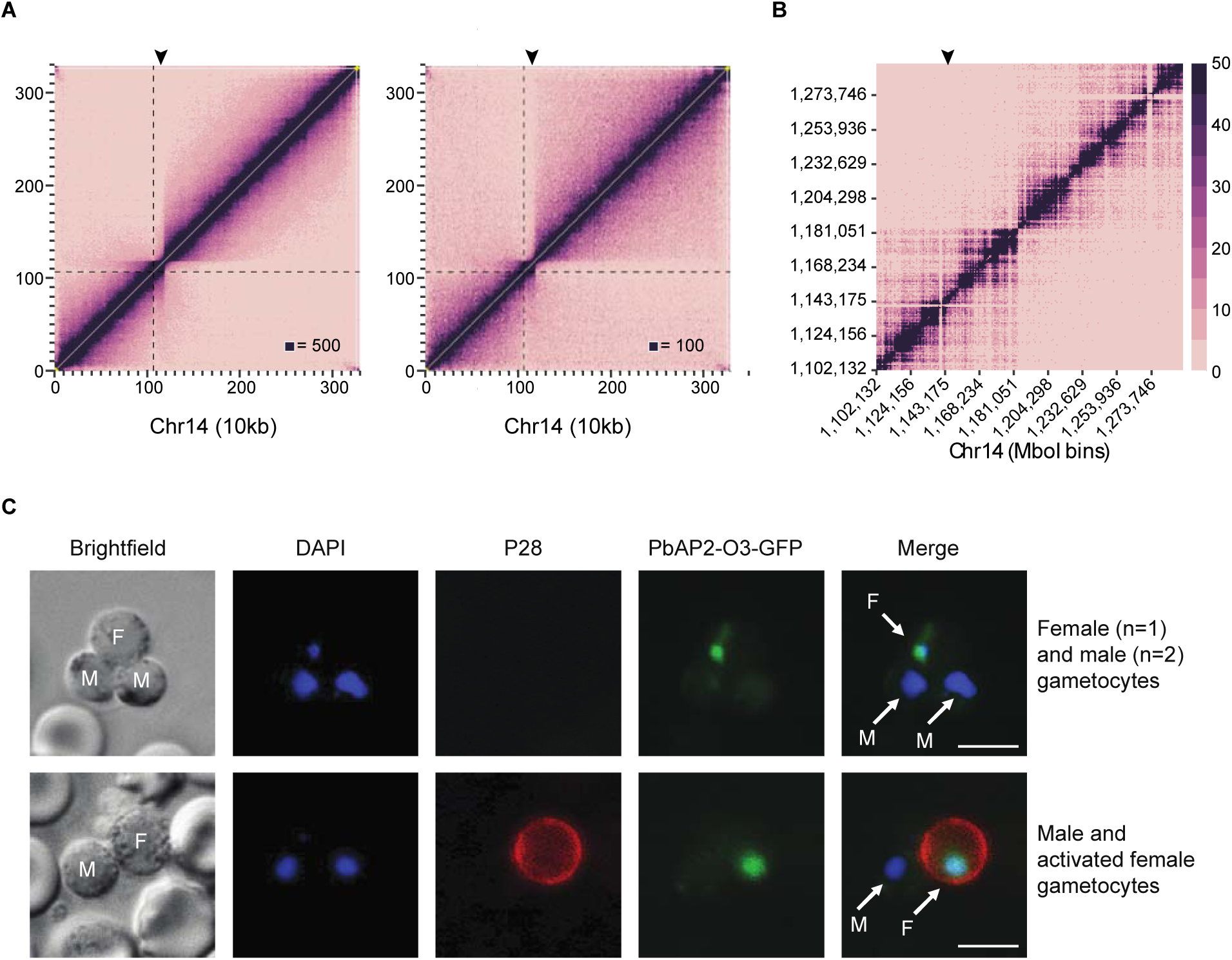
Formation of superdomains on chr14 in gametocytes. **(A)** ICE-normalized contact count heatmap at 1G kb resolution of early gametocyte (left) and late gametocyte (right) chromosome 14 showing the separation of the chromosome into two superdomains. The dashed line indicates the location of the centromere, and the arrowhead indicates the position of PF3D7_14292GG. **(B)** Smaller region of chromosome 14 centered on the domain boundary thatis located inside PF3D7_1430000, a conserved gene with unknown function. **(C)** The homolog of *pfap2* gene PF3D7_1429200 in *P. berghei* (PBANKA_1015500; *pbap2-o3)* has a nuclear localization in female gametocytes and gametes, but is not detected in male gametocytes. The top row shows male and female gametocytes. The bottom row shows a male and female gamete activated by mosquito ingestion, which triggers expression of the female-specific surface protein P28. Male **(M)** and female **(F)** parasite are indicated in the brightfield and merged images. Scale bar denotes 10 μm.

In humans, serine/threonine protein phosphatase 2A (PP2A) is one of the four major Ser/Thr phosphatases and is thought to play a complex, but mostly inhibitory role in the control of cell growth and division^47^. Gametocytes have a higher expression level of *pfptpa* than the IDC stages and express a different variant of *pfptpa* that does not contain exon 1 (**Supplemental Fig. S15A**). The sequence of intron 1 is unusual and contains many motifs that are repetitive (for example, 12 repeats of motif TGTACATACACTTAT and minor variations thereof, within the 705 nt intron; **Supplemental Fig. S15B**). These could be the binding sites for a lncRNA or protein involved in formation of the domain boundary.

The domain boundary is also relatively close to the pfap2-encoding locus PF3D7_1429200 (chr14:1,144,518–1,148,078) (**Supplemental Fig. S14A**). To evaluate whether this TF could be involved in sexual differentiation, we generated a transgenic *P. berghei* strain in which the homolog of PF3D7_1429200 (PBANKA_1015500) was expressed as a GFP-tagged protein (**Supplemental Fig. S14B-C**). *P. berghei* is widely used as a model for *P. falciparum*, in part because of the higher efficiency of genetic manipulations as compared to *P. falciparum*. Female gametocytes, activated female gametes, and zygotes all expressed the tagged ApiAP2 TF with nuclear localization of the protein, while the protein was completely absent in male gametocytes and gametes (**Figure 4C**), as well as in IDC stages (data not shown). These results demonstrate that this ApiAP2 TF (named PfAP2-O3 hereafter in line with a recent publication^48^) is expressed in a strict sex-specific fashion. Additional experiments will be necessary to demonstrate that the formation of the domain boundary is involved in the expression of *pfap2-o3* and thus plays a role in the differentiation of female gametocytes.

### Rearrangement of chromosomes in sporozoites

Similar to the re-localization of invasion genes during the transition from the IDC to the gametocyte stage, the invasion genes also interacted more strongly with virulence genes in *P. falciparum* sporozoites than during the IDC (**Figure 2E** and **Supplemental Fig. S8C**). In *P. vivax* sporozoites, a cluster of invasion genes on chromosome 10 showed depletion of interactions with other loci on the same chromosome as compared to surrounding genomic regions (**Supplemental Fig. S16B**). This observation may suggest that the invasion genes also have a distinct genome organization at this stage of the *P. vivax* life cycle. However, these results may also be caused by sequence variation in invasion genes in the field isolates used in this study as compared to the reference genome, resulting in lower mapping to this region.

In addition, distinct changes were noticeable around rDNA loci. *P. falciparum* encodes four rDNA units containing single copies of the 28S, 5.8S, and 18S genes. The units on chromosomes 5 and 7 are active during the human blood stages, whereas the units on chromosomes 1 and 13 are active in the mosquito stages. Several additional rDNA genes are located on other chromosomes, including a unit of three 5S genes on chromosome 14 (**Figure 5A**). In general, sporozoites showed a large increase in the number of contacts between rDNA genes and virulence genes as compared to the IDC stages and gametocytes (**Figure 5B** and **Supplemental Fig. S8D**). These changes in conformation were most visible in chromosome 7, in which the rDNA unit is located at the boundary of two large domains in the IDC stages and gametocytes, which presumably contributes to its activation status. In sporozoites on the other hand, the separation of the chromosome into two large domains disappeared (**Figure 5C** and **Supplemental Fig. S4B**). Similar changes in domain conformation can be observed around the rDNA locus on chromosome 5 (**Supplemental Fig. S8E**). Unfortunately, the *P. vivax* rDNA genes have not been annotated and we therefore cannot validate whether these genes have a similar impact on genome organization as observed in *P. falciparum*.

**Figure 5:**
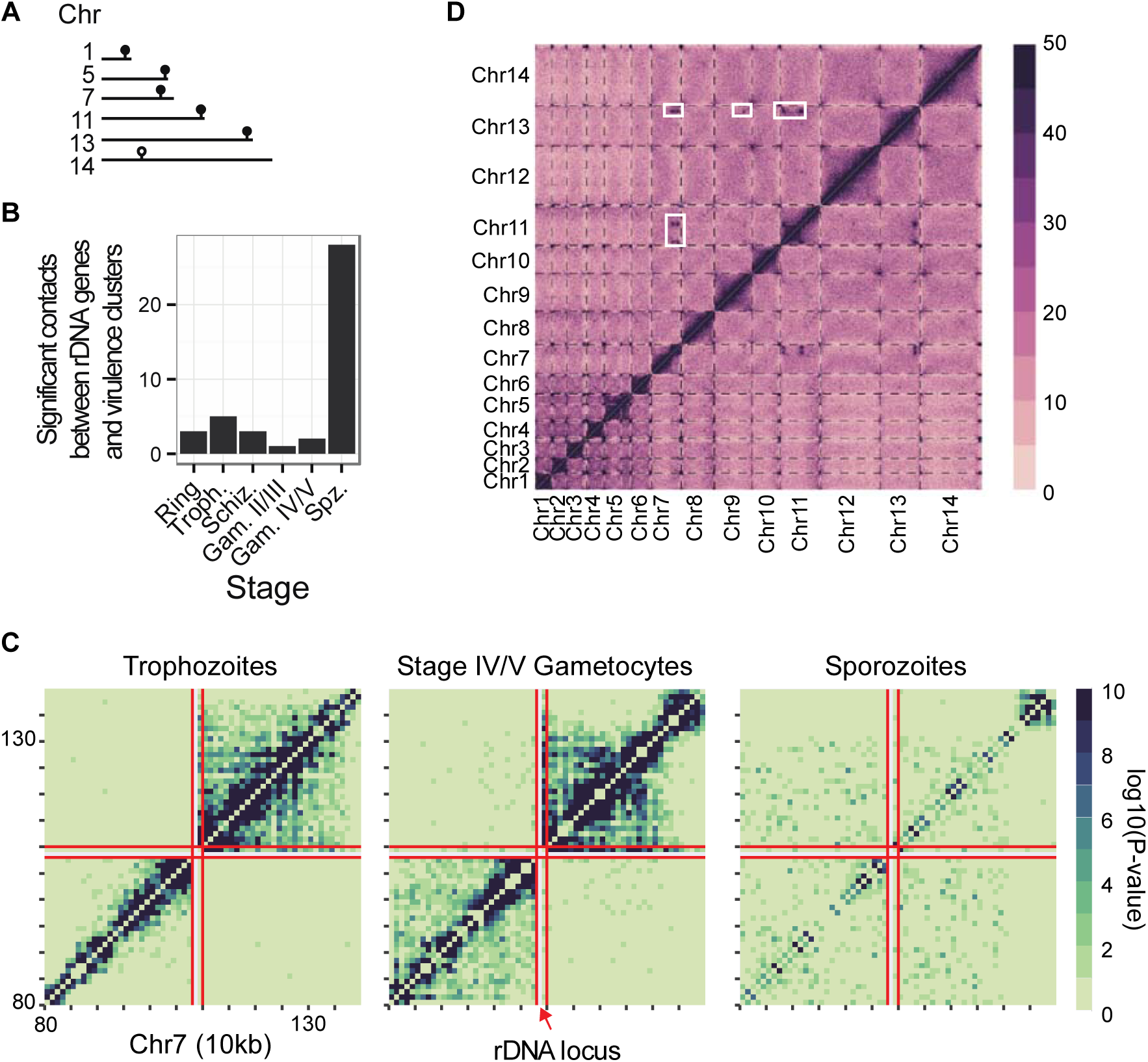
Changes in genome organization in salivary gland sporozoites. **(A)** Locations of rDNA genes in the *P. falciparum* genome. Units of 28S, 5.8S, and 18S genes are indicated with a filled symbol; the unit of three 5S genes is indicated with an open symbol. **(B)** Increased overall number of interactions between rDNA genes and virulence genes in *P. falciparum* sporozoites. **(C)** Loss of domain formation around the rDNA locus on chr7 in *P. falciparum* sporozoites as compared to other life cycle stages. The borders of the rDNA locus are indicated by red lines. **(D)** Strong interchromosomal interactions in *P. vivax* sporozoites, indicated by white rectangles. Dashed lines indicate chromosome boundaries.

Prominent features of genome organization in *P. falciparum* and *P. vivax* sporozoites were strong long-range and interchromosomal contacts that involved other genes than virulence genes and that did not seem to be present in the IDC or gametocyte stages. In *P. vivax*, strong intrachromosomal contacts were present in chromosomes 7 and 11, which also formed strong interchromosomal interactions with each other and with additional loci on chromosomes 9 and 13 (**Figure 1B**, rightmost panel, **Figure 5D**, and **Supplemental Table S7**). For *P. falciparum*, intrachromosomal interactions were observed in chromosomes 3, 4, 8, 9, 11, 13, and 14 (**Supplemental File S1**), although, interestingly, these are not homologous to the loci that participate in loops in *P. vivax*. Several of these loops involved *pfap2* loci and genes involved in sporozoite migration to the liver and in some cases in hepatocyte invasion, such as circumsporozoite protein (PfCSP), sporozoite micronemal protein essential for cell traversal (PfPLP1), thrombospondin-related anonymous protein (PfTRAP), sporozoite protein essential for cell traversal (PfSPECT1), and gamete egress and sporozoite traversal protein (PfGEST) (Table 1).

**Table 1:** Loci involved in long-range intrachromosomal interactions in *P. falciparum* sporozoites.

| Chr | locus (kb)<sup>a</sup> | gene | description | Pv homolog |
| --- | --- | --- | --- | --- |
| 3 | 115 | PF3D7_0302100 | Ser/Thr protein kinase 1 (PfSTPK1) | PVX_119250 |
| 3 | 225 | PF3D7_0304600 | CS protein (PfCSP) | PVX_119355 |
| 4 | 245 | PF3D7_0404500 | 6-cys protein (P52) | PVX_001020 |
| 4 | 375 | PF3D7_0407600 | Conserved, unknown function | n.a. |
| 4 | 425 | PF3D7_0408700 | Perforin -Like Protein 1 (PfPLP1) | PVX_000810 |
| 8 | 135 | PF3D7_0801900 | Conserved, unknown function | PVX_093645 |
| 8 | 295 | PF3D7_0805200<br>PF3D7_0805300 | Gamete release protein (PfGAMER)<br>Conserved, unknown function | PVX_093500<br>PVX_093495 |
| 9 | 125<br>135 | PF3D7_0902800<br>PF3D7_0902900<br>PF3D7_0903000<br>PF3D7_0903100<br>PF3D7_0903200 | Serine repeat antigen 9 (PfSERA9)<br>Conserved, unknown function<br>Conserved, unknown function<br>PfRER1<br>PfRAB7 | N.a.<br>N.a.<br>PVX_098595<br>PVX_098600<br>PVX_098605 |
| 9 | 325 | PF3D7_0906600 | Zinc finger protein | PVX_098775 |
| 9 | 535<br>545 | PF3D7_0911700<br>PF3D7_0911800<br>PF3D7_0911900<br>PF3D7_0912000<br>PF3D7_0912100 | GTP-binding protein<br>Conserved, unknown function<br>Falstatin (PfICP)<br>Conserved, unknown function<br>Zinc finger protein | PVX_099025<br>PVX_099030<br>PVX_099035<br>PVX_099040<br>PVX_099045 |
| 11 | 225 | PF3D7_1105000<br>PF3D7_1105100<br>PF3D7_1105200 | Histone H4 (PfH4)<br>Histone H2B (PfH2B)<br>WD repeat-containing protein (PfWRAP73) | PVX_090930<br>PVX_090935<br>PVX_090940 |
| 11 | 335 | PF3D7_1107800 | ApiAP2 TF | PVX_091065 |
| 13 | 1,465 | PF3D7_1335900 | PfTRAP | PVX_082740 |
| 13 | 1,675 | PF3D7_1342500 | PfSPECT1 | PVX_083025 |
| 14 | 1,875 | PF3D7_1445600<br>PF3D7_1445700 | RNA-binding protein<br>Conserved, unknown function | PVX_118205<br>PVX_118200 |
| 14 | 2,005 | PF3D7_1449000 | PfGEST | PVX_118040 |

Kb, kilobase; Pv, *Plasmodium vivax*; n.a., not available.
<sup>a</sup> All loci listed interact with all other listed loci in the same chromosome.

### Identification of a putative clonally variant gene family in P. vivax

The genome annotation of *P. vivax* is less complete than that of *P. falciparum*, and many genes have been grouped into families based solely on sequence homology. An example is the *Pv-fam-e* (also named *rad)* gene family that is closely related to the *Plasmodium* helical interspersed sub-telomeric *(phist*) gene family^49^, encoding exported proteins involved in erythrocyte remodeling^50–52^. The *P. vivax* genome contains 45 *rad* genes, of which 10 and 27, respectively, are located in two separate clusters on chromosome 5. In the contact count matrix of *P. vivax* chromosome 5, the largest of the two clusters strongly interacted with the (sub-)telomeric regions and showed a depletion of interactions with all other intrachromosomal loci (**Supplemental Fig. S16C**). These results suggest that *P. vivax rad* genes may be regulated by organization into facultative heterochromatin. Additional experiments, such as H3K9me3 ChIP-seq analysis, will be necessary to demonstrate whether this is indeed the case.

### PfHP1 is essential for virulence gene colocalization

The clustering of virulence genes seems to be a general feature of the *P. falciparum* genome that is maintained throughout its life cycle. Recently, it was shown that depletion of PfHP1 results in loss of *var* gene repression and an arrest in parasite growth^35^, suggesting that this protein is essential for structural integrity of the repressive cluster. To study the effect of PfHP1 depletion on genome conformation, we performed Hi-C experiments on a transgenic *P. falciparum* strain expressing PfHP1 fused to GFP and a destabilization domain (DD), both in the presence and in the absence of Shield-1, resulting in expression or knockdown of the PfHP1 fusion protein, respectively^35^. Ring-stage parasites expressing the tagged PfHP1 protein showed a decrease in virulence gene clustering as compared to wild-type parasites (p-value = 0.026 and p-value < = 0.001, respectively, BH FDR-corrected Paulsen colocalization test^53^ (see **Computational Methods** in **Supplemental Information**). In particular, interactions between internal and subtelomeric *var* gene clusters were lost (**Supplemental Fig. S17**). This result is in agreement with an increase in internal *var* gene expression in this strain, as reported previously^35^. Virulence gene clustering was completely lost in the PfHP1-depleted strain (p-value = 0.129), in line with a generalized loss of *var* gene repression^35^. Accordingly, we observed more significantly changing virulence gene bins as compared with wild-type in the PfHP1-depleted strain (n=71) than in the PfHP1-tagged strain (n=18). These results confirm that PfHP1 is indeed essential for maintenance of the structure of the repressive cluster and thus for regulation of virulence gene expression.

### The 3D genome structure correlates with gene expression

Finally, we explored the relationship between gene expression and 3D structure, leveraging four published expression data sets^54–57^ and our 3D models of the genome architecture. As in our previous study, we applied kernel canonical correlation analysis (KCCA)^58^. KCCA is an unsupervised learning approach akin to principal component analysis that identifies a set of orthogonal gene expression components that are coherent with the 3D structure. To perform this analysis, we separated the 3D models into three distinct groups: those related to IDC (ring, trophozoite, schizont), gametocyte (early and late) and sporozoite stages. For each of these groups of time points, we extracted a gene expression component and a structure component that exhibited coherence to the expression profiles and 3D structure, such that genes whose expression is correlated with the selected gene expression component tend to be colocalized in 3D. The gene expression components for the three sets of structures were highly correlated and were dominated by the repressive center (**Figure 6A** and ref. 20). To further interpret the results of the KCCA, we extracted ranked lists of genes based on their KCCA scores: lists that rank genes based on the similarity of their gene expression profiles with the first or second gene expression components, and lists that rank genes based on the similarity of their 3D position with the first or second structure components. We then investigated whether several sets of genes were enriched in those ranked lists. Var, *rif*, and exported protein genes all showed strong and significant enrichment on the first gene expression and structure components for all stages (**Figure 6B**), as expected based on their localization in or near the repressive center. In contrast, the invasion genes were significantly enriched for the first gene expression component in gametocytes and sporozoites, and for the second gene expression and structure components in all stages (**Figure 6B**). While the average scores for this group of genes are relatively small, these results suggest that expression of these genes is also coordinated with their location within the nucleus. Gametocyte-specific genes did not show correlation to the first or second component for either gene expression or structure, which is in line with regulation of these genes by other factors, such as the gametocyte-specific transcription factor PfAP2-G, instead of localization within the nucleus.

**Figure 6:**
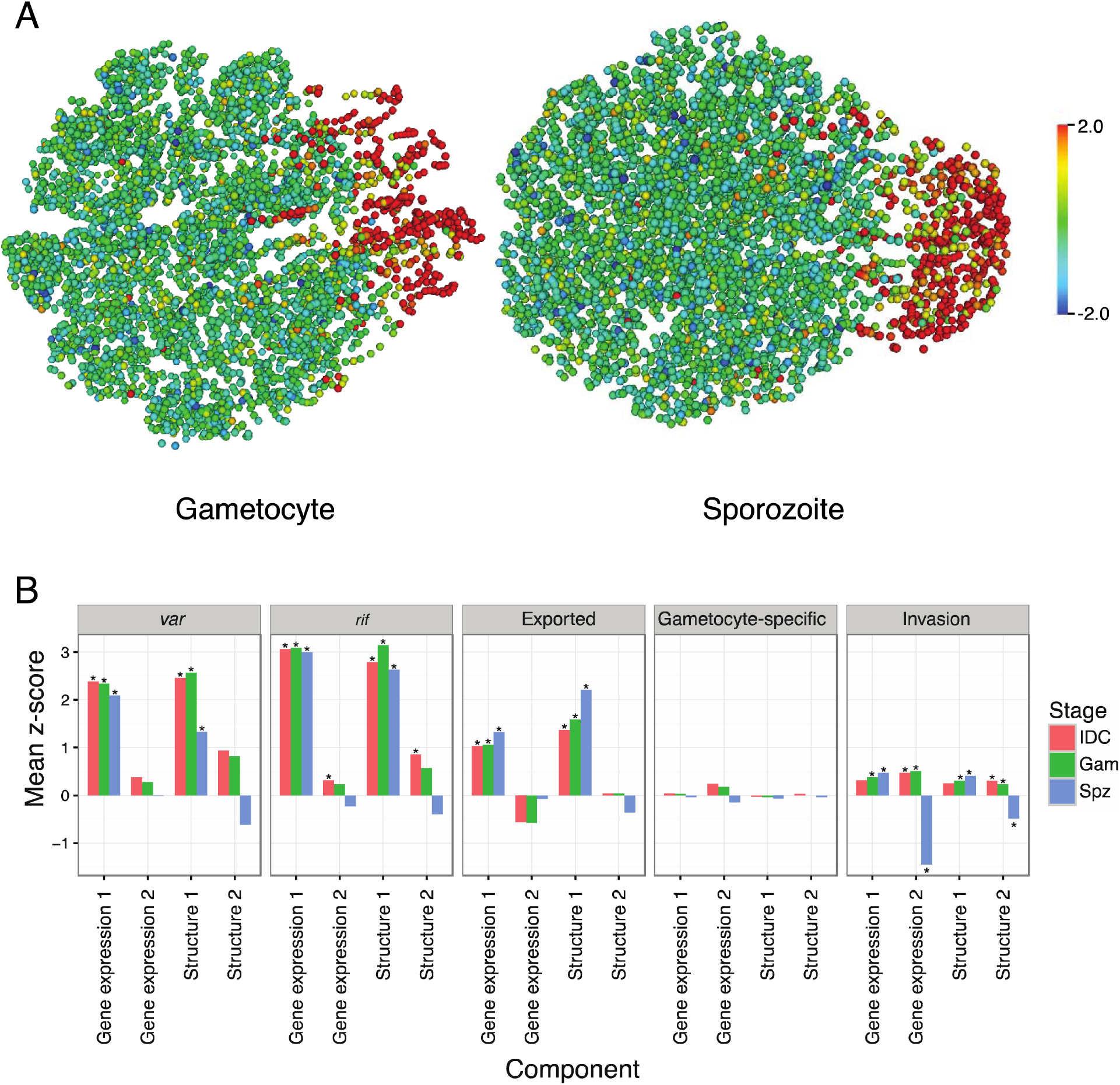
3D genome structure correlates with gene expression. **(A)** Each gene is plotted by its position within the 3D structure and is colored by its standardized KCCA score (see **Computational Methods**) in the first gene expression component. The direction of this first gene expression component is from the telomere cluster on the right to the opposite side of the nucleus and is dominated by the repressive center (genes colored in red). **(B)** The average standardized KCCA score for the first and second gene expression components and the first and second structure components of specific groups of genes. Stars indicate groups of genes for which the standardized KCCA scores for both the gene expression component and the structure component were significantly different (t-test, FDR < 0.1%).

## DISCUSSION

Understanding the mechanisms involved in gene regulation during the various life cycle stages of *P. falciparum* will be important for the development of novel strategies to block parasite replication and transmission. Our previous study described the hallmarks of genome organization during the intra-erythrocytic developmental stages of *P. falciparum*, and showed that nuclear architecture correlated well with gene expression^20^. Here, we investigated chromosome conformation and chromatin structure in the stages of parasite transmission from human to mosquito (gametocytes) and from mosquito to human (sporozoites) and compared all stages to identify various subsets of genes that exhibit changes in genome organization during the complex life cycle of *Plasmodium* parasites.

Our results confirm that virulence gene families, such as *var, rifin*, and *stevor*, cluster in perinuclear heterochromatin, not only during the IDC stages, but also in gametocytes and sporozoites. Previous reports showed multiple virulence gene clusters scattered around the nucleus, either by DNA-FISH on telomere repeat sequences^16,17^, IFA on H3K9me3^9^ or IFA on proteins that bind to chromosome ends (PfSIP2-N^59^) or subtelomeric regions (PfHP1^35,59^). Our population-based Hi-C data showed strong interactions between all telomeres, which could be consistent with a completely random distribution of telomeres over multiple clusters. However, our IFA results conflict with this large body of data and instead show a single H3K9me3 focus in all blood stages in which the parasite is not undergoing DNA replication, representative of a single repressive center. Similar results were previously obtained by IFA against the heterochromatin mark H3K36me3^15^ and also recently for heterochromatin mark H3K9me3^60^. Differences in timing during the cell cycle (before or after start of DNA replication) may account for some of these differences. Indeed, we also observed multiple H3K9me3 foci during the trophozoite stage, most likely as a result of nuclear expansion and the start of DNA multiplication. We currently do not have an explanation that would reconcile our observations with those of others in the field. However, whether *var* genes cluster in one or multiple repressive centers throughout the IDC, both models are consistent with the presence of distinct heterochromatin and euchromatin regions controlling gene expression and antigenic variation in *P. falciparum*.

In gametocytes, the overall genome organization is similar to IDC stages, in line with relatively small changes in the gene expression program during gametocytogenesis^61^. We observed specific changes for *pfap2-g*^5^ that dissociates from the repressive center, and for invasion genes that associate with the repressive center. In addition, subtelomeric genes encoding exported proteins are selectively silenced or activated by H3K9me3 deposition or removal, respectively. These results confirm that remodeling of the infected host cell is among the essential changes that occur during gametocytogenesis^62,63^, and that an epigenetic switch provides an extra layer of transcriptional regulation for the genes involved. The presence of H3K9me3 in the relatively large intergenic region downstream of *pfgdv1* suggests that this region by itself is an important determinant in the regulation of gametocyte differentiation. A lncRNA is transcribed from this region at low levels during the IDC^64^, and it is tempting to speculate that this transcript is upregulated in gametocytes and possibly essential for gametocyte development.

The sharp division of chromosome 14 into two superdomains is an intriguing finding that will need to be explored in more detail. The inactivated X chromosome (Xi) in mammalian cells adopts a seemingly similar structure with a ~200 kb hinge region. However, the domain boundary on *P. falciparum* chromosome 14 is sharp and does not have a distinct hinge region. In addition, the domain boundary on Xi is organized into euchromatin, while the surrounding chromosome is in a heterochromatic state^65^. In contrast, we did not observe a distinct pattern in H3K9me3 marking around the domain boundary on chromosome 14. If the large domain boundary in *P. falciparum* has a similar function as that on Xi, this would suggest that boundary formation may only occur in one of the two sexes and that paternal or maternal heritage of chromosome 14 may influence gene expression during or after sexual replication in the zygote, ookinete, or oocyst stage.

In support of this hypothesis, we discovered that the homolog of ApiAP2 transcription factor PF3D7_1429200 (PfAP2-O3) in *P. berghei* is strictly female-specific, suggesting that PfAP2-O3 controls female gametocyte differentiation. A few hundred genes are differentially expressed between male and female gametocytes^66,67^ and could be under the control of this protein, either through direct transcriptional regulation or through selective stabilization of female-specific transcripts. Interestingly, disruption of *ap2-o3* in *P. berghei* resulted in differential gene expression in gametocytes and reduced gametocyte levels, although the sex ratio was not influenced^48^. However, *pfap2-o3* is located approximately 40 kb from the domain boundary and it is not directly clear if and how expression of this gene would be influenced by the formation of these superdomains.

An alternative hypothesis is that the domain boundary is involved in regulation of the nearby gene Pf3D7_1430100 *(pfptpa)*. PfPTPA has been shown to bind and activate PP2A, and to block the G2/M transition^68^. The transition from rapidly dividing asexual parasites into cell cycle arrested gametocytes is likely to require tight cell cycle regulation by a protein such as PTPA. Given the differences in *pfptpa* expression between the IDC and gametocyte stages, we speculate that the domain boundary inside or close to this gene may be important for driving the expression of this gene, which may in turn activate PP2A to block cell division in gametocytes. In ongoing efforts, we are further investigating the role of the domain boundary in gene expression and gametocyte differentiation.

From an evolutionary perspective, *P. falciparum* and *P. vivax* are highly divergent within the family of *Plasmodium* species^69,70^. It is therefore striking that both species show similar organization of their genomes in the sporozoite stage, in particular a large reduction in centromere clustering and numerous long-range interactions, including several strong interchromosomal interactions in *P. vivax*. The salivary gland sporozoite maintains a relatively quiescent transcriptional state, waiting for injection into the human bloodstream and invasion of a hepatocyte before ramping up transcriptional activity. Our interpretation of the genome organization in the sporozoite stage is that the majority of the genome is transcriptionally repressed, with the exception of several active loci that colocalize in transcriptional islands, giving rise to long-range interactions. The observation that the genes involved in these long-range interactions are not each other’s homologs in *P. falciparum* and *P. vivax* is suggestive of species-specific gene expression and warrants further investigation.

To fully understand transcriptional regulation, it is of great interest to unravel the causal relationship between genome organization and transcriptional activity. In multicellular organisms, evidence is accumulating that certain aspects of genome organization are independent of transcription^71^. In addition, disruptions in genome structure that bring together previously isolated promoters and enhancers can result in gene activation^72^. It will be important to determine to what extent genome organization controls transcriptional activity in *P. falciparum*. Our findings bring a new level of insight into genome dynamics during the *Plasmodium* life cycle and open up new avenues for targeted approaches towards understanding parasite gene regulation. In addition, molecules inhibiting the (re-)structuring of the genome have the potential to act as potent transmission-blocking antimalarials.

## METHODS

### Experimental procedures

#### Parasite strains and cultures

The *P. falciparum* strain NF54 was cultured in human O* erythrocytes at 5% haematocrit as previously described^73^. The induction of gametocyte-stage parasites was adapted from a previously published protocol^74^. Stage IV/V gametocytes were isolated using a percoll gradient, were cultured for one additional day, and were then isolated by magnetic purification yielding 6.25 × 10^8^ parasites (**Supplemental Fig. S1**). Adult female *Anopheles stephensi* mosquitoes were allowed to feed on *P. falciparum* NF54 gametocyte cultures. Sporozoites were harvested from infected mosquitoes 14 to 19 days later.

Stage II/III gametocytes were obtained using the *P. falciparum* NF54^Ps16^ reporter gene line^75^ Gametocytes were isolated by magnetic purification, yielding 1.17 × 10^8^ parasites with high purity (>95% gametocytes) as determined by GFP expression (**Supplemental Fig. S1**). To obtain *P. vivax* sporozoites, female *Anopheles cracens* mosquitoes were fed on blood samples drawn from *P. vivax* infected patients who had signed a consent form (protocol approved by Oxford Tropical Research Ethics Committee) and who attended a Shoklo Malaria Research Unit (SMRU) clinic in Mawker Thai or Wang Pha, on the western Thailand-Myanmar border. Fifteen days post-infection, *P. vivax* sporozoites were harvested from *An. cracens* salivary glands. The construction of *P. falciparum* 3D7 transgenic strain PfHP1-GFP-DD has been described previously^35^. Parasites were synchronized, split into two populations at 4–12 hpi and cultured in the presence or absence of Shield-1 as described. Parasites were harvested for Hi-C at 4–12 hpi in the next cell cycle. More details about parasite cultures and collections are provided in **Supplemental Information**.

#### Hi-C procedure

Parasites were crosslinked in 1.25% formaldehyde for 25 min as previously described^20^ in a total volume between 1 and 10 ml, depending on the number of parasites harvested. For late stage gametocytes, the crosslinking protocol was slightly modified. Late gametocytes were collected in lysis buffer (25 mM Tris-HCl, pH 8.0, 10 mM NaCl, 2 mM 4-(2-aminoethyl)benzenesulfonyl fluoride HCl (AEBSF), 1% Igepal CA-360 (v/v), and EDTA-free protease inhibitor cocktail (Roche)) and incubated for 10 min at RT. After homogenization by 15 needle passages, formaldehyde was added to a final concentration of 1.25%, followed by 10 more needle passages. The protocol was then continued as for all other samples. To map the inter- and intrachromosomal contact counts, crosslinked parasites were subjected to the tethered conformation capture procedure as previously described^20,76^ with minor modifications, using MboI for restriction digests. Details are provided in **Supplemental Information**.

#### DNA-FISH

DNA-FISH experiments were performed as previously described^20^. Sequences of primers used for probe generation are shown in **Supplemental Table S8**.

#### Immunofluorescence microscopy

*P. falciparum* IDC-stage parasites and gametocytes were fixed onto slides using 4% paraformaldehyde for 30 min at RT. Slides were washed three times using 1x PBS. The parasites were permeabilized with 0.1% Triton-X for 30 min at RT, followed by a wash step with 1x PBS. Samples were blocked overnight at 4°C in IFA buffer (2% BSA, 0.05% Tween-20, 100 mM glycine, 3 mM EDTA, 150 mM NaCl and 1x PBS). Cells were incubated with anti-Histone H3 antibody (ab8898 (Abcam), 1:500 or 07–442 (Millipore), 1:500) for 1 hr at RT followed by anti-rabbit Alexa Fluor 488 (Life Technologies A11008; 1:500). No differences were observed in the results obtained with the two primary antibodies (**Supplemental Fig. S11**). Slides were mounted in Vectashield mounting medium with DAPI. Images were acquired using an Olympus BX40 epifluorescence microscope.

#### H3K9me3 ChIP-seq

Asexual parasites were crosslinked for 10 min with 1% formaldehyde in PBS at 37°C, while gametocytes were crosslinked with 1.25% formaldehyde in lysis buffer for 25 min at RT. Chromatin was sheared using the Covaris Ultra Sonicator (S220) ChIP was performed as described previously^77^ using 2 μg of anti-Histone H3K9me3 antibody (ab8898 (Abcam) for biological replicates #1 and 07–442 (Millipore) for biological replicates #2) or no antibody as a negative control. Details are provided in **Supplemental Information**.

#### Validation of sex-specific ApiAP2 TF in P. berghei

Transgenic parasites endogenously expressing a GFP-tagged version of the *P. berghei* homolog of Pf3D7_1429200 (PBANKA_1015500) were generated by single homologous recombination^78^. For the endogenously C-terminal fusion GFP-tagged parasites, a diagnostic PCR reaction was used as illustrated in **Supplemental Fig. S14C**. Phenotypic analyses were performed as previously described^78,79^. Details are provided in **Supplemental Information**.

### Computational methods

We mapped and binned reads and created and ICE-normalized contact count matrices as previously described (^20^ and **Supplemental Information**). To identify significant contacts, we modeled the effect of genomic distance on contact count probability with a spline using fit-hi-c^23^. Because of the possibility of differing statistical power between datasets, we report both the number of significant contacts above a given threshold, as well as the percent of significant contacts meeting a criterion (e.g. between *pfap2* gene loci and virulence clusters) out of all significant contacts. Significant co-localization of gene sets was assessed using previously developed tests (^25,80^ and **Supplemental Information**).

We developed ACCOST (Altered Chromatin Conformation STatistics) to estimate the statistical significance of differences in contact counts between samples, taking as inspiration the negative binomial-based tests used for RNA-seq data^27,81,82^. We adapted the model employed by DESeq to Hi-C data by using an explicit specific scaling factor corresponding to bin-specific ICE biases. In addition, we estimated variance and dispersion of the negative binomial without replicates by assuming that most bins at a given genomic distance act similarly. For details of the model and implementation, see **Supplemental Information**.

## DATA AVAILABILITY

The sequencing data from this study have been submitted to the NCBI Sequence Read Archive under accession numbers SRP091967 (Hi-C) and SRP091939 (ChIP-seq). Fold-change heatmaps can be accessed at http://noble.gs.washington.edu/proj/plasmo3dsexualstages/. The source code for ACCOST is available at https://github.com/cookkate/ACCOST.

## ACKNOWLEDGEMENTS

We thank Till Voss (Swiss Tropical and Public Health Institute) for preparation of the PfHP1 transgenic strain samples, Clay Clark, John Weger, and Glenn Hicks (Institute for Integrative Genome Biology, University of California Riverside) for their assistance in Illumina sequencing, and Xueqing (Maggie) Lu for help with ChIP-seq analysis. We are grateful to the Insectary and Parasitology Core Facilities at the Johns Hopkins Malaria Research Institute and in particular would like to thank Abai Tripathi, Godfree Mlambo and Chris Kizito for their outstanding work and The Bloomberg Family Foundation for supporting these facilities. SMRU is part of the Mahidol-Oxford University Research Unit, supported by the Wellcome Trust. The following reagents were obtained through the MR4 as part of the BEI Resources Repository, NIAID, NIH: NF54 (MRA-1000) deposited by Megan Dowler, Walter Reed Army Institute of Research. This study was financially supported by the National Institutes of Health (grants R01 AI85077–01A1 to KGLR, R01 AI06775–01 to WSN and KGLR, and R01 AI056840 to PS), the University of California, Riverside (NIFA-Hatch-225935 to KGLR), the European Research Council (grant ERC-SMAC-280032 to NV and JPV), the French National Research Agency (grant ABS4NGS ANR-11-BINF-0001 to NV and JPV), the Miller Institute for Basic Research in Science (JPV), the Fulbright Foundation (JPV), the Bill & Melinda Gates Foundation (OPP1040938 to DAF), the Medical Research Council UK (grant MR/K011782/1 to RT), the Human Frontier Science Program (postdoctoral fellowship LT000507/2011-L to EMB), the University of Texas Health Science Center at San Antonio (EMB), the Natural Sciences and Engineering Research Council of Canada (postdoctoral fellowship to KBC), the Gordon and Betty Moore Foundation (Grant GBMF3834 to UC Berkeley), the Alfred P. Sloan Foundation (Grant 2013–10–27 to UC Berkeley), Institute Leadership Funds from La Jolla Institute for Allergy and Immunology (FA), and used the Extreme Science and Engineering Discovery Environment (XSEDE), which is supported by National Science Foundation grant number ACI-1548562.

## AUTHOR CONTRIBUTIONS

WSN and KGLR conceived the project. EMB and GB carried out the Hi-C and ChIP-seq experiments. JP maintained *P. falciparum* cultures and assisted in experimental procedures. LS, CA, LSR, DAF, FN, and PS prepared parasite samples for Hi-C experiments. DB and RT performed experiments in *P. berghei*. KBC and NV analyzed the data under the supervision of WSN and JV. FA contributed to data analysis. EMB, KBC, NV, FA, WSN, and KGLR wrote the manuscript. All authors reviewed the manuscript.

## MATERIALS & CORRESPONDENCE

Correspondence and material requests should be addressed to Karine G. Le Roch or William S. Noble.

